# Somatic Mutations in *MCOLN3* in Aldosterone-Producing Adenomas cause Primary Aldosteronism

**DOI:** 10.1101/2024.10.20.619295

**Authors:** Desmaré van Rooyen, Sascha Bandulik, Grace Coon, Miriam Laukemper, Chandan Kumar-Sinha, Aaron M. Udager, Chaelin Lee, Heather Wachtel, Debbie L. Cohen, James M. Luther, Thomas Giordano, Adina Turcu, Richard Warth, William E. Rainey, Juilee Rege

**Author notes:** co-first authors (Desmaré van Rooyen, Sascha Bandulik). **Corresponding author**: Juilee Rege **Email:**. **Competing Interest Statement:** James M. Luther (consulting for Mineralys).

## Abstract

Primary aldosteronism is characterized by renin-independent hyperaldosteronism that originates from aldosterone-producing lesions in the adrenal glands. Under physiological conditions, aldosterone synthase (*CYP11B2*) expression is confined to the adrenal zona glomerulosa where it catalyzes the final reaction yielding aldosterone. The regulation of *CYP11B2* transcription depends on the control of cellular membrane potential and cytosolic calcium activity. In primary aldosteronism, aldosterone-producing adenomas (APAs) are characterized by disrupted regulation of CYP11B2 expression resulting in autonomous biosynthesis of aldosterone. These lesions often harbor aldosterone-driver somatic mutations in genes encoding ion transporters/channels/pumps that increase cytosolic calcium activity causing increased *CYP11B2* expression and aldosterone biosynthesis. We investigated APAs devoid of known somatic mutations and detected a missense mutation and a deletion-insertion variant in *MCOLN3* which encodes for mucolipin-3 (TRPML3) — a highly conserved inwardly-rectifying, cation-permeable channel. These *MCOLN3* mutations were identified in three APAs derived from male patients with primary aldosteronism: p. Y391D and p.N411_V412delinsI. Both mutations are located near the ion pore and selectivity filter of TRPML3. This is the first report of disease-causing *MCOLN3* mutations in humans. Functional studies suggest *MCOLN3^Y391D^* might directly or indirectly via membrane depolarization alter calcium influx of transfected adrenocortical cells, resulting in increased *CYP11B2* transcription and aldosterone production. This study implicates mutated *MCOLN3* as a driver of aldosterone excess in primary aldosteronism.

**Significance Statement:** Primary aldosteronism is a common but under-diagnosed endocrine disease that contributes to global hypertension burden and cardiovascular mortality and morbidity. Hyperaldosteronism in primary aldosteronism is mainly caused by adrenal lesions harboring somatic mutations that disrupt intracellular calcium levels and consequently aldosterone synthase expression and aldosterone production. Majority of these mutations have been identified in genes encoding ion transporters/channels/pumps. Herein, we report the first disease-causing somatic mutations in human *MCOLN3* in aldosterone-producing adenomas (APAs) devoid of known mutations. *In vitro* investigations showed the *MCOLN3* variant (p.Y391D) caused an influx of cytosolic calcium in adrenocortical cells and the subsequent increase in aldosterone synthase and aldosterone biosynthesis.

## Introduction

Primary aldosteronism is the most prevalent cause of secondary hypertension, accounting for 5-15 % (1–4) and 14-21 % (5) of the general- and resistant hypertension cases, respectively. Primary aldosteronism is characterized by renin-independent dysregulated production of aldosterone from one (unilateral) or both (bilateral) adrenal glands resulting in chronic hypertension.

Aldosterone excess in unilateral primary aldosteronism is most often attributed to lesions defined by the recent consensus on the histopathology of primary aldosteronism (HISTALDO) as aldosterone-producing adenomas (APAs) or, to smaller lesions called aldosterone-producing nodules (APNs) (6). Autonomous aldosterone biosynthesis in these lesions is caused by aldosterone-driver somatic mutations in genes which mainly translate to cell surface ion pumps, transporters and channels. These mutations induce cytosolic calcium influx through cell membrane depolarization, and in turn, upregulate aldosterone synthase (CYP11B2) expression and aldosterone production. Next-generation sequencing (NGS) has identified aldosterone-driver mutations in genes including *KCNJ5* (potassium channel) (7–9), *ATP1A1* (sodium/potassium ATPase) and *ATP2B3* (plasma membrane calcium ATPase) (10, 11), *CACNA1D* (10, 12, 13) and *CACNA1H* (14) (voltage-gated calcium L- and T-type channels, respectively), *CLCN2* (chloride channel) (15, 16), and most recently *SLC30A1* (zinc transporter) (17). Mutations have also been found in genes encoding proteins other than ion transporters including *CADM1* (18), *CTNNB1* (13, 19), *PRKACA* (20), *and GNAQ/11* (21).

The advances in adrenal lesion DNA-capture guided by CYP11B2 immunohistochemistry (IHC) and sequencing technologies have improved the detection rate of somatic mutations to 89-98 % (8, 14, 22). This, in turn, has provided important clues into the pathogenesis of primary aldosteronism. Our sequencing pipeline led to the identification of novel aldosterone-driver mutations in *MCOLN3* that encodes mucolipin-3/TRPML3 (transient receptor potential cation channel, 3). *MCOLN3* was deemed to be a potential cause of primary aldosteronism since autonomous aldosterone production in this disease has been shown to be a consequence of dysregulated ion conductance (23). Herein, we describe the discovery and characterization of new *MCOLN3* variants. *MCOLN3* has mostly been studied in context of autophagy (24–28) with little known regarding its role in the adrenal gland. Importantly, *MCOLN3* mutations have not been previously reported in APAs or APNs nor has this gene been associated with disease in humans. In this study, we have demonstrated that novel *MCOLN3* mutations result in increased intracellular calcium that leads to elevated adrenal cell aldosterone production.

## Results

We investigated the differential expression of *MCOLN3* mRNA in multiple normal human tissues and found highest expression in the adrenal cortex (Fig. 1A). As compared to RNA isolated from adrenal, *MCOLN3* levels were found to be significantly higher in APAs (2-fold; *P*<0.05) while no significant difference was observed in cortisol-producing adenomas (CPAs) (Fig. 1B). *CYP11B2* mRNA expression was significantly higher in APAs (102-fold; *P*<0.001) compared to the adrenal, while *CYP11B2* expression in CPAs was expectedly low (Fig. 1B).

**Figure 1.**
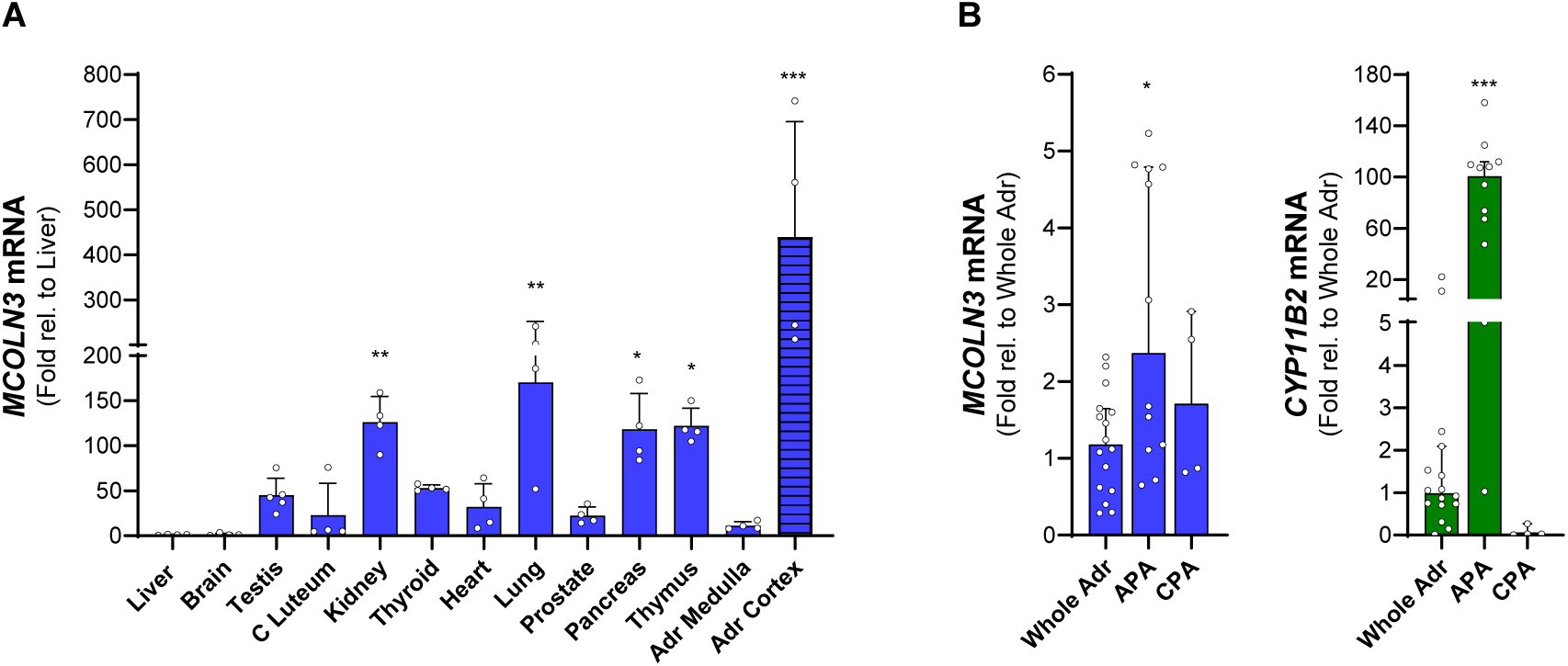
*MCOLN3* mRNA expression in human tissue and adrenocortical adenomas. Real-time quantitative PCR (qRT-PCR) analyses of (**A**) *MCOLN3* mRNA in multiple snap frozen human tissues showed highest expression in the adrenal cortex. *MCOLN3* expression is expressed and compared relative to liver *MCOLN3* expression (**B**) *MCOLN3* and *CYP11B2* mRNA analysis of adrenocortical-derived tissues expressed and compared relative to the whole adrenal. Aldosterone-producing adenomas (APAs), but not cortisol-producing adenomas (CPAs), exhibited significantly higher *MCOLN3* expression compared to the adrenal. As opposed to the whole adrenal, *CYP11B2* transcript levels in APAs, but not CPAs, were increased. Data is presented as the (**A**) mean ±SD or (**B**) median with 95% confidence interval of at least four tissue samples normalized to *PPIA*. Significance was analyzed using a Kruskal-Wallis with a Dunn’s multiple comparison test. \**P*<0.05, \*\**P*<0.01, \*\*\**P*<0.001.

Previous investigations of morphologically normal adrenals reported the age-related change in expression pattern of CYP11B2 highlighting that not all ZG cells are aldosterone-producing. Continuous zona glomerulosa CYP11B2 expression was associated more with younger adrenals while older adrenals presented with discontinuous or no CYP11B2 expression in the zona glomerulosa (29). We, therefore, also investigated the protein expression of MCOLN3 in adrenal glands with continuous zona glomerulosa CYP11B2 expression, and in adrenal glands with a discontinuous CYP11B2-positive zona glomerulosa with aldosterone-producing micronodules (APMs), respectively (representative adrenals in Fig. 2). Results showed that immunoreactivity for MCOLN3 was localized to the zona glomerulosa in both tissue types. MCOLN3 and CYP11B2 expression co-localized in the zona glomerulosa of the adrenal with classical adrenocortical CYP11B2-positive zona glomerulosa architecture. However, while CYP11B2 expression in the adrenal with APMs was sporadic, MCOLN3 expression remained continuous throughout the CYP11B2-negative zona glomerulosa.

**Figure 2.**
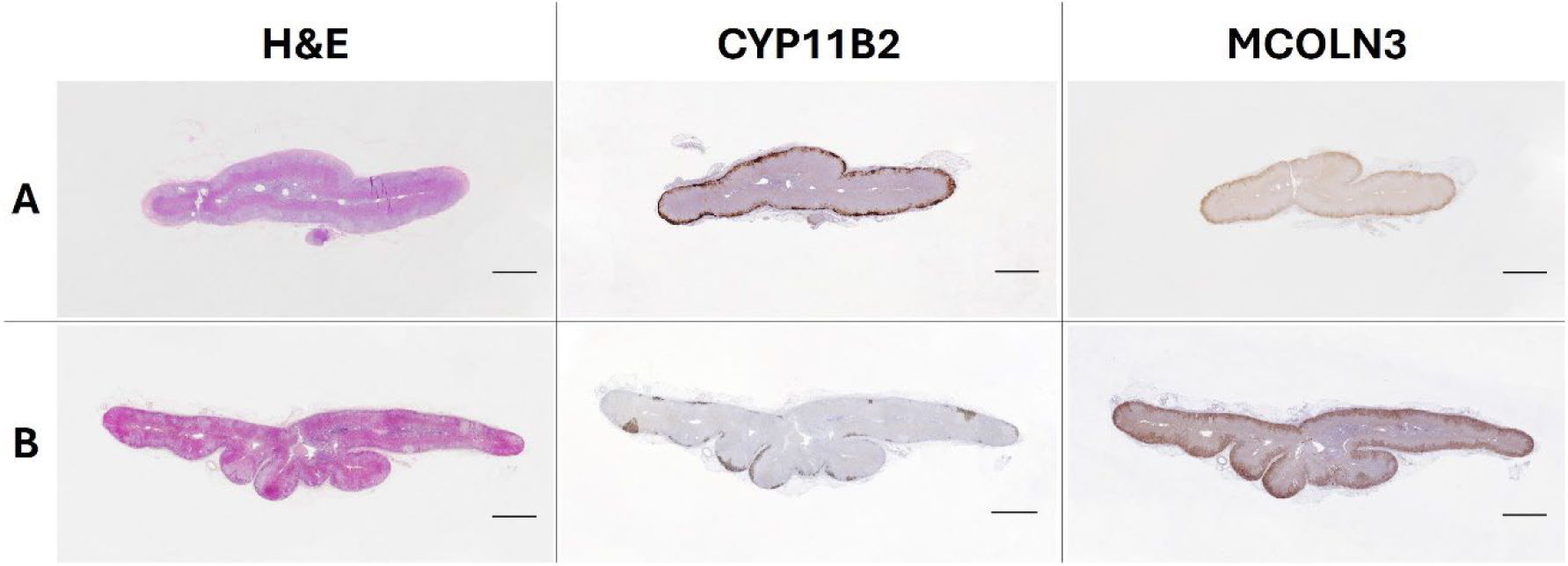
Histological characteristics of human adrenals from healthy subjects. Hematoxylin and eosin (H&E) staining, aldosterone synthase (CYP11B2) and mucolipin- 3 (MCOLN3) immunohistochemistry staining of 5 µm FFPE sections representative of (**A**) an adrenal with classical adrenal architecture and (**B**) an adrenal with disrupted zona glomerulosa CYP11B2 expression with aldosterone-producing micronodules (APMs). Scale bar, 2 mm.

CYP11B2 IHC-guided whole exome sequencing (WES) analysis of two formalin-fixed paraffin-embedded (FFPE) APA tissues devoid of known mutations and their matched adjacent adrenal tissue identified a novel somatic mutation in *MCOLN3*. This missense mutation p.Y391D (NM_018298.11: c.1171 T>G) was confirmed by Sanger sequencing (Fig. 3A). We, therefore, updated our targeted NGS panel to include *MCOLN3* and discovered an additional APA with an in-frame deletion-insertion in *MCOLN3* resulting in the loss of asparagine and valine at positions 411 and 412, respectively, and the insertion of isoleucine (p.N411_V412delinsI; *MCOLN3^N411_V412delinsI^*) at this position. The *MCOLN3^N411_V412delinsI^* was also validated by Sanger sequencing (Fig. 3B). These mutations are in close proximity to the MCOLN3 pore and ion selectivity filter (Fig. 3C). Furthermore, Tyr391 and Val412 are highly conserved in orthologs of various species (Fig. 3D) including the mouse in which disease-causing mutations in *MCOLN3* has been reported (30–32).

**Figure 3.**
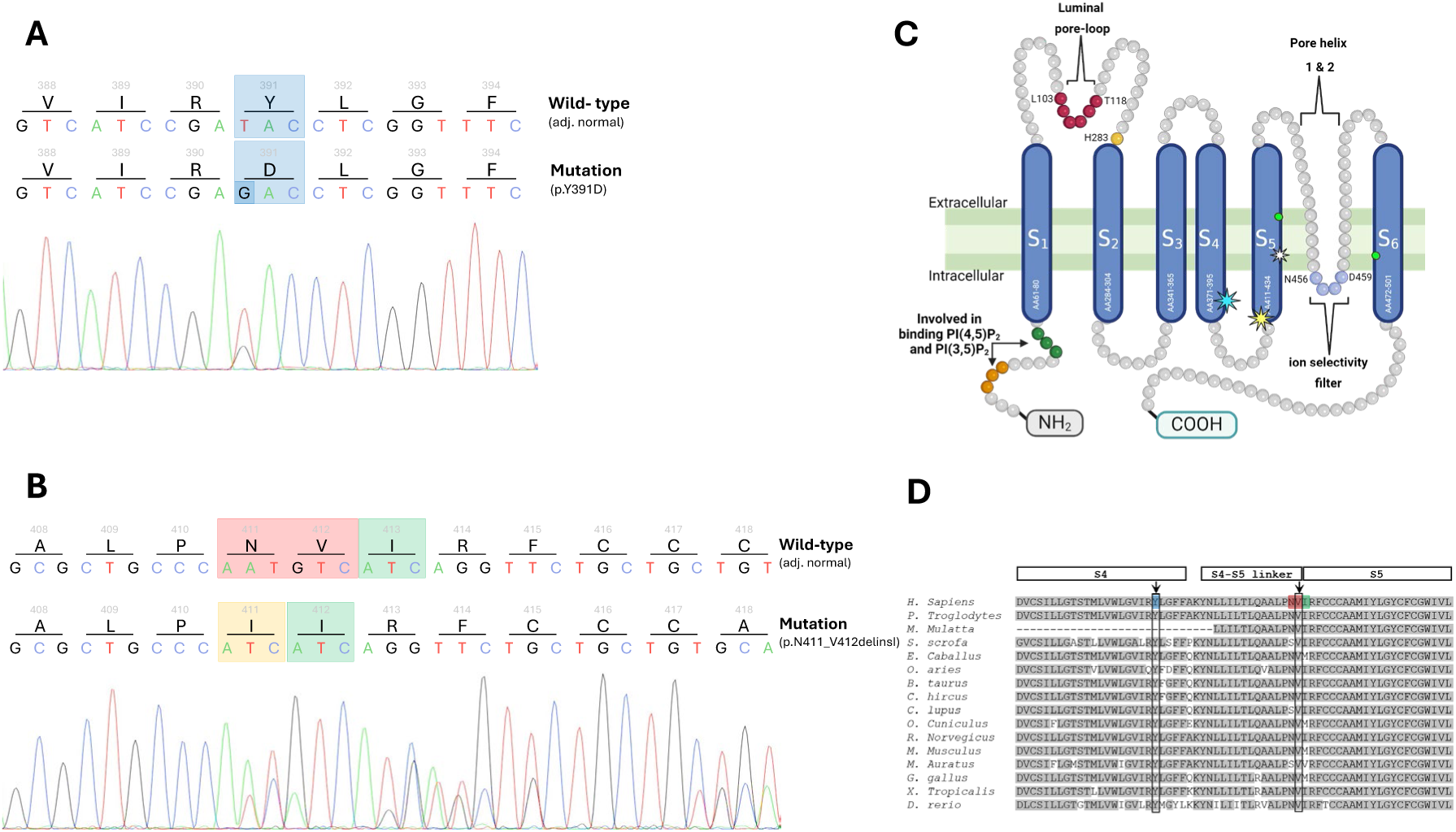
*MCOLN3* mutations in APAs. Sanger sequence chromatograms of tumor and adjacent adrenal tissue DNA for validation of the *MCOLN3* variants. **A)** Translational consequence of a point mutation in the *MCOLN3^Y391D^* variant resulting in the single residue change, Tyr391Asp (blue). **B)** *MCOLN3^N411_V412deIinsI^*variant presents with the deletion of two amino acids, Asn411 and Val412 (red), and the insertion of Ile411(yellow) causing a single residue frame shift (green). Mutations are absent in the tumor adjacent adrenal tissue (top sequence). **C)** Simplified schematic representation of the MCOLN3 transmembrane adapted from the work done by Zhou et al. (56). The location of *MCOLN3^Y391D^*(cyan star) and *MCOLN3^N411_V412deIinsI^* (yellow star) in segment 4 (S4) and segment 5 (S5), respectively, is shown in close proximity to the ion selectivity filter. The disease-causing *Mcoln3^A419P^* variant identified in mice is shown in S5 by the white star. Neon green circles mark the amino acids involved in binding of MCOLN3 agonist, ML-SA1. **D)** Alignment of orthologs among various species shows the conservation of Tyr391 and Val412 in the mucolipin-3 protein. Affected amino acid residue associated with the novel variants are highlighted in blue (*MCOLN3^Y391D^*) and red and green (*MCOLN3^N411_V412deIinsI^*). Highlighted in grey are residues conserved in orthologs.

Collectively, *MCOLN3* mutations were identified in three APAs, all derived from male patients with primary aldosteronism across different racial groups between the ages of 57-76 years (Table 1). Cross-sectional imaging and adrenal vein sampling (AVS) confirmed that two patients (patients ELA25 and AA30) presented with a solitary aldosterone-producing adrenal tumor. The third patient (patient UM102), however, exhibited with an aldosterone-producing lesion on the left adrenal gland and a non-functional adenoma on the contralateral gland. While the aldosterone-to-renin ratio (ARR) was severely elevated in all three patients with primary aldosteronism (178 to <1000), only 2 of 3 patients presented with moderate hypokalemia. The patient harboring the *MCOLN3^N411_V412delinsI^* presented with more severe hypertension and hypokalemia compared to the patients with the missense mutation. However, this patient was on fewer antihypertensive medications and of African ancestry which might suggest a higher sensitivity to sodium and aldosterone and consequently more aggravated symptoms (33–35).

**Table 1.**
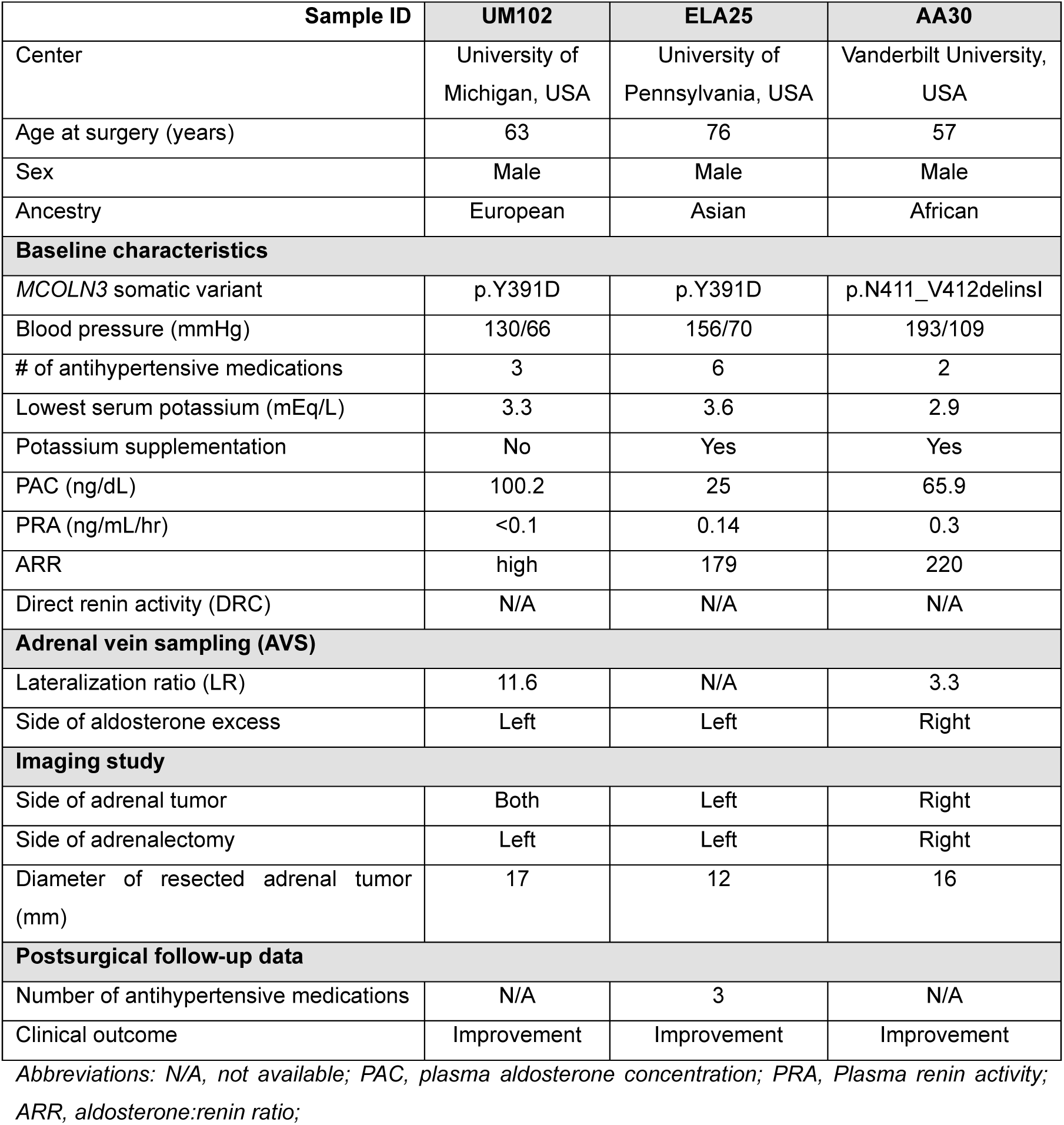
Clinical characteristics of patients with APAs harboring somatic *MCOLN3* mutations.

| Sample ID | UM102 | ELA25 | AA30 |
| --- | --- | --- | --- |
| Center | University of Michigan, USA | University of Pennsylvania, USA | Vanderbilt University, USA |
| Age at surgery (years) | 63 | 76 | 57 |
| Sex | Male | Male | Male |
| Ancestry | European | Asian | African |
| **Baseline characteristics** |  |  |  |
| *MCOLN3* somatic variant | p.Y391D | p.Y391D | p.N411_V412delinsI |
| Blood pressure (mmHg) | 130/66 | 156/70 | 193/109 |
| # of antihypertensive medications | 3 | 6 | 2 |
| Lowest serum potassium (mEq/L) | 3.3 | 3.6 | 2.9 |
| Potassium supplementation | No | Yes | Yes |
| PAC (ng/dL) | 100.2 | 25 | 65.9 |
| PRA (ng/mL/hr) | <0.1 | 0.14 | 0.3 |
| ARR | high | 179 | 220 |
| Direct renin activity (DRC) | N/A | N/A | N/A |
| **Adrenal vein sampling (AVS)** |  |  |  |
| Lateralization ratio (LR) | 11.6 | N/A | 3.3 |
| Side of aldosterone excess | Left | Left | Right |
| **Imaging study** |  |  |  |
| Side of adrenal tumor | Both | Left | Right |
| Side of adrenalectomy | Left | Left | Right |
| Diameter of resected adrenal tumor (mm) | 17 | 12 | 16 |
| **Postsurgical follow-up data** |  |  |  |
| Number of antihypertensive medications | N/A | 3 | N/A |
| Clinical outcome | Improvement | Improvement | Improvement |

Abbreviations: N/A, not available; PAC, plasma aldosterone concentration; PRA, Plasma renin activity; ARR, aldosterone:renin ratio;

Histologic assessment of mutated *MCOLN3*-harboring tumors (Fig. 4) showed increased immunoreactivity for MCOLN3 and CYP11B2 in tumors with the missense mutation as compared to the deletion-insertion variant. MCOLN3 expression in the APA with the *MCOLN3^N411_V412delinsI^* variant was sparse as compared to the zona glomerulosa of adjacent adrenal tissue. In contrast, adjacent adrenal tissue of the APA harboring the *MCOLN3^N411_V412delinsI^* variant was devoid of CYP11B2 except for a single APM while CYP11B2 immunoreactivity in the tumor was heterogenous. CYP17A1 expression was also observed in the tumors with the *MCOLN3^Y391D^* mutation but less prominent in the *_MCOLN3_N411_V412delinsI* _variant._

**Figure 4.**
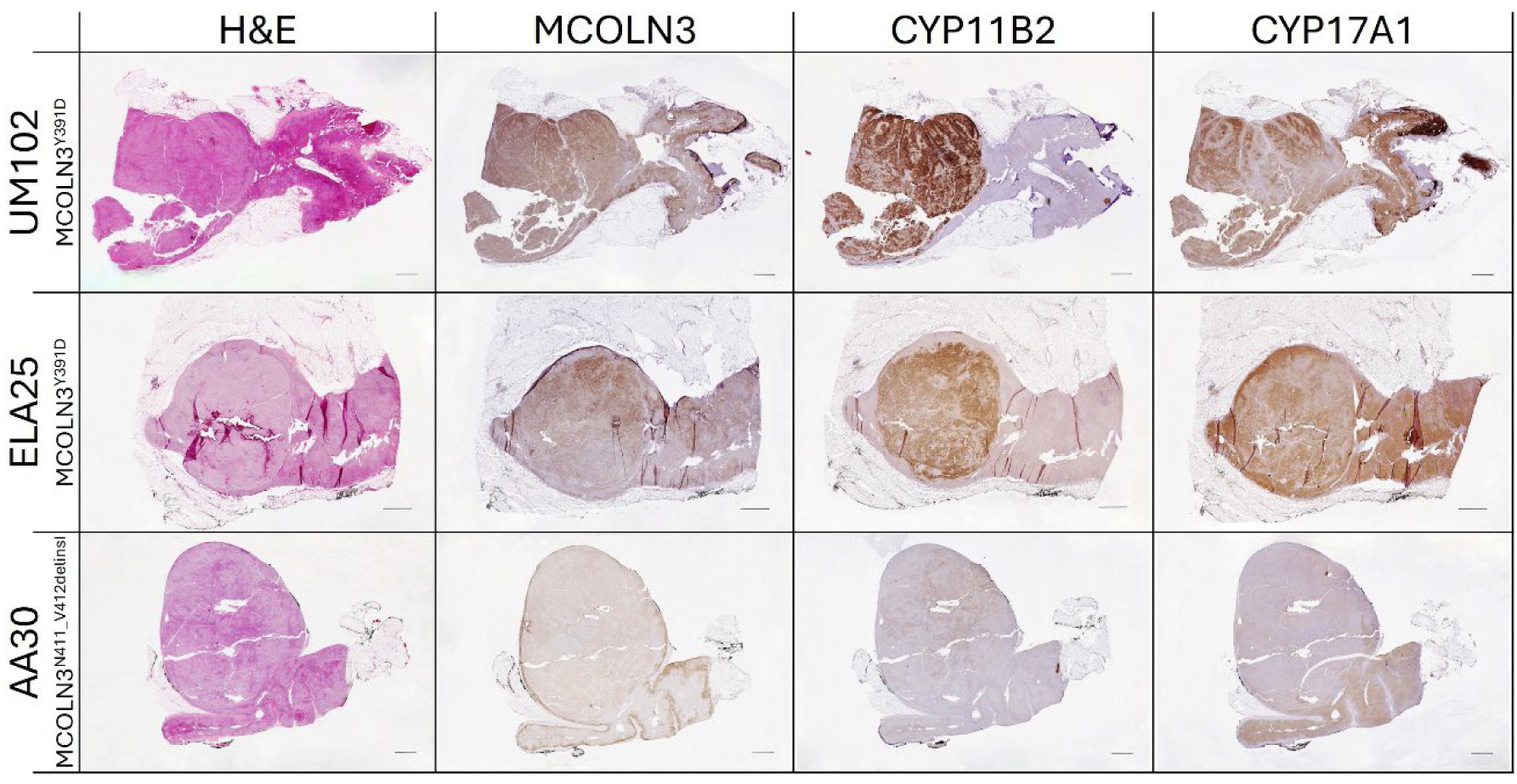
Histopathological characteristics of the APAs with somatic MCOLN3 mutations. Hematoxylin and eosin (H&E) staining, and MCOLN3, CYP11B2 and CYP17A1 immunohistochemistry staining of the resected APAs identified with the *MCOLN3^Y391D^* (UM102 and ELA25) and *MCOLN3^N411_V412delinsI^* (AA30) variants. Scale bar, 2 mm.

To characterize the functional effects of the *MCOLN3^Y391D^*variant that was identified in 2 of the 3 APAs, we transfected HAC15 adrenocortical cells with plasmid DNA encoding *MCOLN3^Y391D^* (HAC15-*MCOLN3^Y391D^*cells) and assessed the changes in *MCOLN3*, *CYP11B2, NR4A2* (NURR1, an upstream regulator of CYP11B2 transcription (36), *CYP11B1* (11β-hydroxylase) and *CYP17A1* (17α-hydroxylase/17,20-lyase) transcript levels and aldosterone production after 48 h. Data was compared to cells transfected with the empty pIRES-CD8 plasmid vector (empty) and the wild type *MCOLN3* variant (HAC15-*MCOLN3^WT^*cells). Transfection was determined to be successful owing to the elevated *MCOLN3* mRNA levels detected in cells transfected with *MCOLN3^WT^* and *MCOLN3^Y391D^* compared to cells transfected with the empty plasmid (*P*<0.0001) (Fig. 5A). No change in transcript levels of *NR4A2*, *CYP11B2, CYP11B1* or *CYP17A1* was observed in cells expressing *MCOLN3^WT^*. In addition, aldosterone production was unaffected in HAC15-*MCOLN3^WT^* cells. In contrast, the role of *MCOLN3^Y391D^* in driving hyperaldosteronism was highlighted by the 14-fold increase in *NR4A2* mRNA (*P*<0.0001) (Fig. 5B) and the 3.5-fold increase in *CYP11B2* mRNA (*P*<0.0001) (Fig. 5C) compared to *MCOLN3^WT^*. Interestingly, while immunoreactivity of CYP17A1 co-localized with MCOLN3 expression in the tumors (Fig. 4), transfected adrenocortical cell experiments showed little change in *CYP17A1* mRNA levels as a result of *MCOLN3^Y391D^* overexpression (Fig. 5D). Although *CYP17A1* mRNA in HAC15-*MCOLN3^Y391D^* cells was slightly increased compared to HAC15-*MCOLN3^WT^* cells (*P*<0.05), no significant change was observed when compared to the empty controls. Inversely, *CYP11B1* expression significantly differed in HAC15-*MCOLN3^Y391D^* cells *vs* HAC15-*MCOLN3^WT^* cells (*P*<0.01) (Fig. 5E). Importantly, a 2.2-fold induction in aldosterone production was observed in the HAC15-*MCOLN3^Y391D^* cells *vs* HAC15-*MCOLN3^WT^* cells. (Fig. 5F). Additionally, the biosynthesis of hybrid steroids, 18-hydroxycortisol and 18-oxocortisol that are associated with lesions co-expressing CYP11B2 and CYP17A1 (37–40) —were found to be elevated in the HAC15-*MCOLN3^Y391D^*cells whereas cortisol remained unchanged (Supplemental Fig. 1).

**Figure 5.**
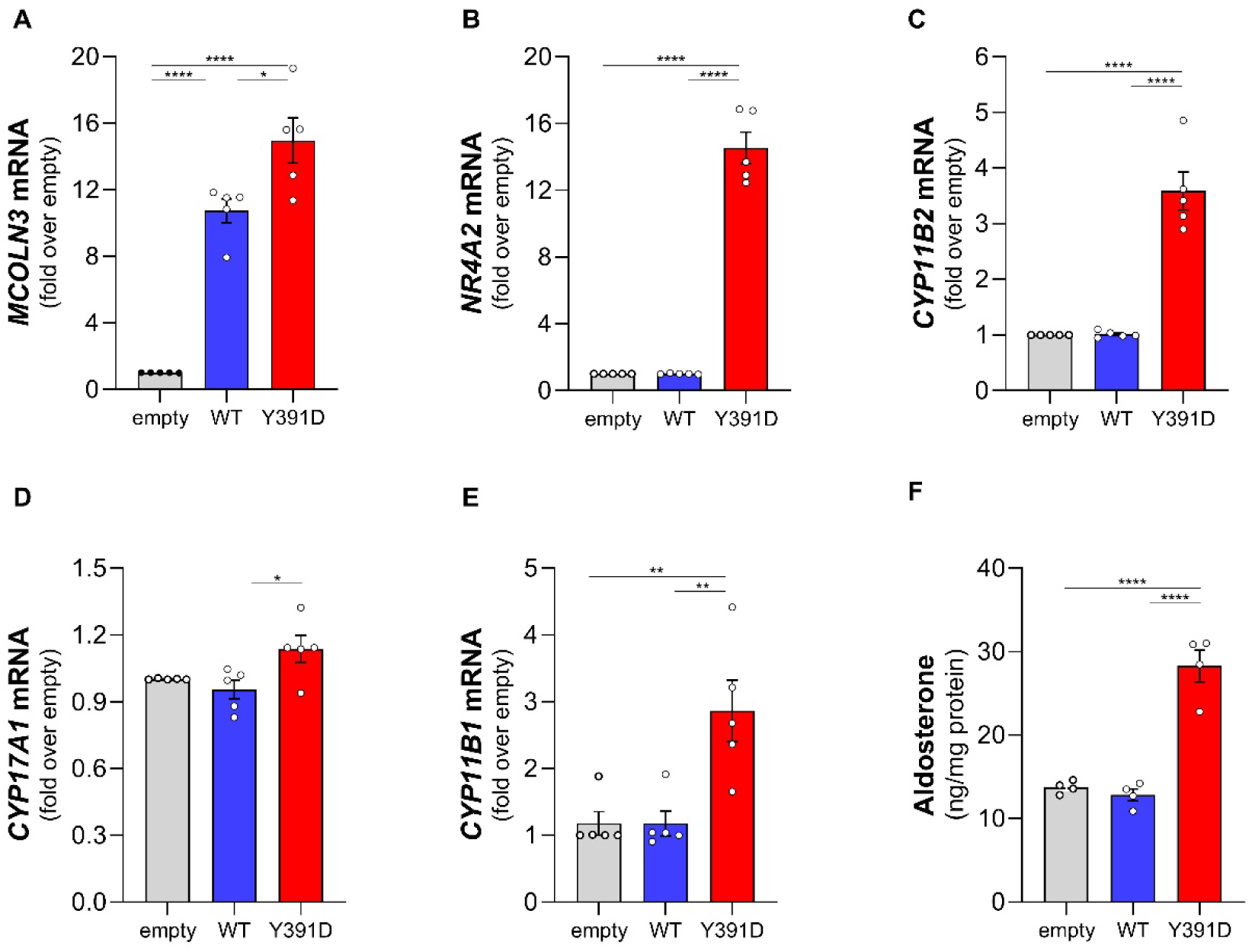
Impact of the *MCOLN3^Y391D^*variant on mRNA expression of *NR4A2*, *CYP11B2*, *CYP17A1* and *CYP11B1*, along with aldosterone production. HAC15 cells were transfected with 1 µg of empty pIRES-CD8 plasmid (empty) or with pIRES-CD8 plasmids containing *MCOLN3^WT^* (WT) or *MCOLN3^Y391D^*(Y391D) via electroporation. 48h post-transfection, the relative mRNA expression levels of (**A**) *MCOLN3*, (**B**) *NR4A2*, (**C**) *CYP11B2*, (**D**) *CYP17A1*, and (**E**) *CYP11B1* were quantified by qRT-PCR. (**F**) Aldosterone content in cell medium was quantified using an ELISA and normalized to total protein content. Data are presented as the mean ±SEM of at least four independent experiments. Each data point represents the mean of the experimental triplicates. Statistical analyses were performed using an ordinary one-way ANOVA with a Tukey’s multiple comparison test. * *P*<0.05; *** *P*<0.001; \*\*\*\**P*<0.0001

Under physiologic conditions, the upregulation of adrenal *CYP11B2* transcription and the consequent production of aldosterone relies on the increase of cytosolic calcium activity.

This can be stimulated by depolarization of the cell membrane, which activates calcium influx through voltage-gated calcium channels, and by calcium release from intracellular calcium stores. Physiologically, depolarization is induced by an increase in extracellular K^+^ and by Ang II-triggered inhibition of K^+^-channels in the plasma membrane. Other primary aldosteronism-associated mutated ion-channels and -transporters cause a pathological depolarization due to an abnormal Na^+^-permeability (7, 17, 41, 42). Since *MCOLN3* is reportedly permeable to calcium and sodium, we tested the ability of *MCOLN3^Y391D^* to similarly cause membrane depolarization and disruption in the regulation of cytosolic calcium levels. We investigated the basal cell membrane potential of HAC15-*MCOLN3^Y391D^* cells by whole-cell patch-clamp recordings and compared it with the empty control and HAC15-*MCOLN3^WT^* cells. The basal membrane potential of the HAC15-*MCOLN3^Y391D^* cells (-13.6 ±2.4 mV) was strongly depolarized compared to the HAC15-*MCOLN3^WT^* (-64.6 ±1.9 mV) and empty control cells (-67.4 ±1.9 mV) (Fig. 6A). To characterize the ion-currents underlying the strong depolarization of HAC15-*MCOLN3^Y391D^* cells, we measured the whole-cell currents at different voltage-clamp steps. Analysis of control, HAC15-*MCOLN3^WT^* and HAC15-*MCOLN3^Y391D^* cells gave outward currents typical of adrenal cell lines and native adrenal glomerulosa cells, which are supposed to be conducted by K^+^ channels that normally hyperpolarize cells (Fig. 7A-C). However, HAC15-*MCOLN3^Y391D^* cells exhibited an additional large inward current at voltages between -100 and -40 mV that was significantly reduced after replacement of extracellular Na^+^ by cell-impermeable NMDG^+^. This was not observed for the empty or HAC15-*MCOLN3^WT^* cells under either condition. In parallel, removing extracellular sodium resulted in a significantly less depolarized cell membrane in HAC15-*MCOLN3^Y391D^* cells (-34.7 ±2.9 mV), albeit cells did not hyperpolarize to the level of control cells. Interestingly, HAC15-*MCOLN3^WT^* cells were also hyperpolarized to some extent by sodium removal (Fig. 7D). Overall, the patch-clamp data suggest an abnormal basal activity of mutant *MCOLN3^Y391D^*, which caused depolarization via increased Na^+^ currents.

**Figure 6.**
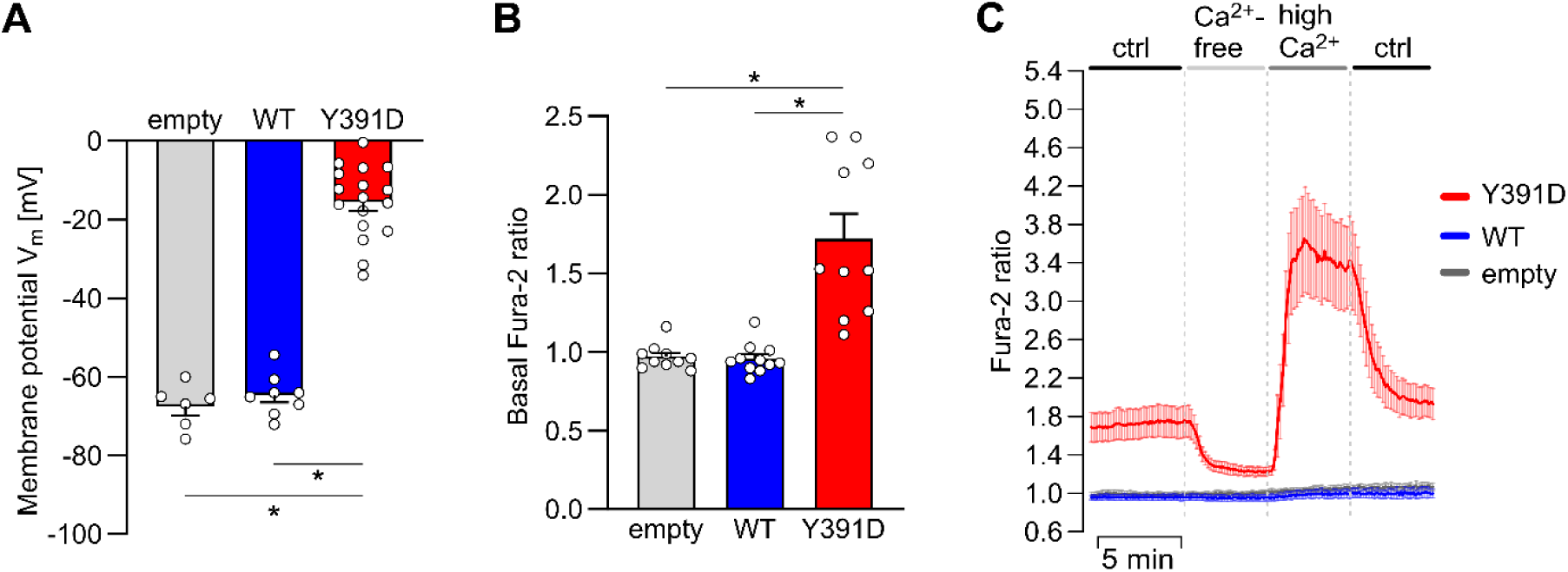
Impact of mutant *MCOLN3^Y391D^* on basal membrane potential and cytosolic calcium (Ca^2+^) activity in transiently transfected adrenal HAC15 cells. HAC15 cells were transfected with 1 µg empty vector (empty) or with 0.5 µg of wild type (WT) or *MCOLN3^Y391D^* (Y391D)-containing plasmids (adjusted to 1 µg plasmid concentration using empty vector). 24-30 h after transfection, cells were used for whole-cell patch-clamp experiments to measure the membrane potential (**A**) and for measurements of the cytosolic calcium activity with fura-2 (**B, C**). (**A**) Membrane potential of cells with mutant *MCOLN3^Y391D^* was depolarized compared to cells with empty vector or with *MCOLN3^WT^* (empty, n=7; *MCOLN3^WT^*, n=8; *MCOLN3^Y391D^*, n=17). (**B**) For calcium measurements, basal calcium activities were calculated from the first control phase of the experiments shown in **C**. Basal calcium activity of cells with *MCOLN3^Y391D^* was increased compared to cells with empty vector or with *MCOLN3^WT^*. Depending on the transfection rate, the signals from 2-6 cells with bound CD8 beads per dish were averaged. Shown here is the mean ±SEM of the mean Fura-2 ratio from n dishes: empty, n=10; *MCOLN3^WT^*, n=11; *MCOLN3^Y391D^*, n=10. Figure **C** summarizes changes of fura-2 ratio over time upon application of the indicated extracellular solutions. Cells with *MCOLN3^Y391D^* showed a greater dependence of cytosolic calcium on extracellular calcium changes. This was characterized by a significant decrease in calcium activity when extracellular calcium was removed and an increase when extracellular calcium was increased. *, *P*<0.05, *MCOLN3^WT^*vs. *MCOLN3^Y391D^* cells or empty CD8. Statistical significance (*) was tested with the GraphPad Prism 10 software using one-way ANOVA analysis with Tukey’s multiple comparison.

**Figure 7.**
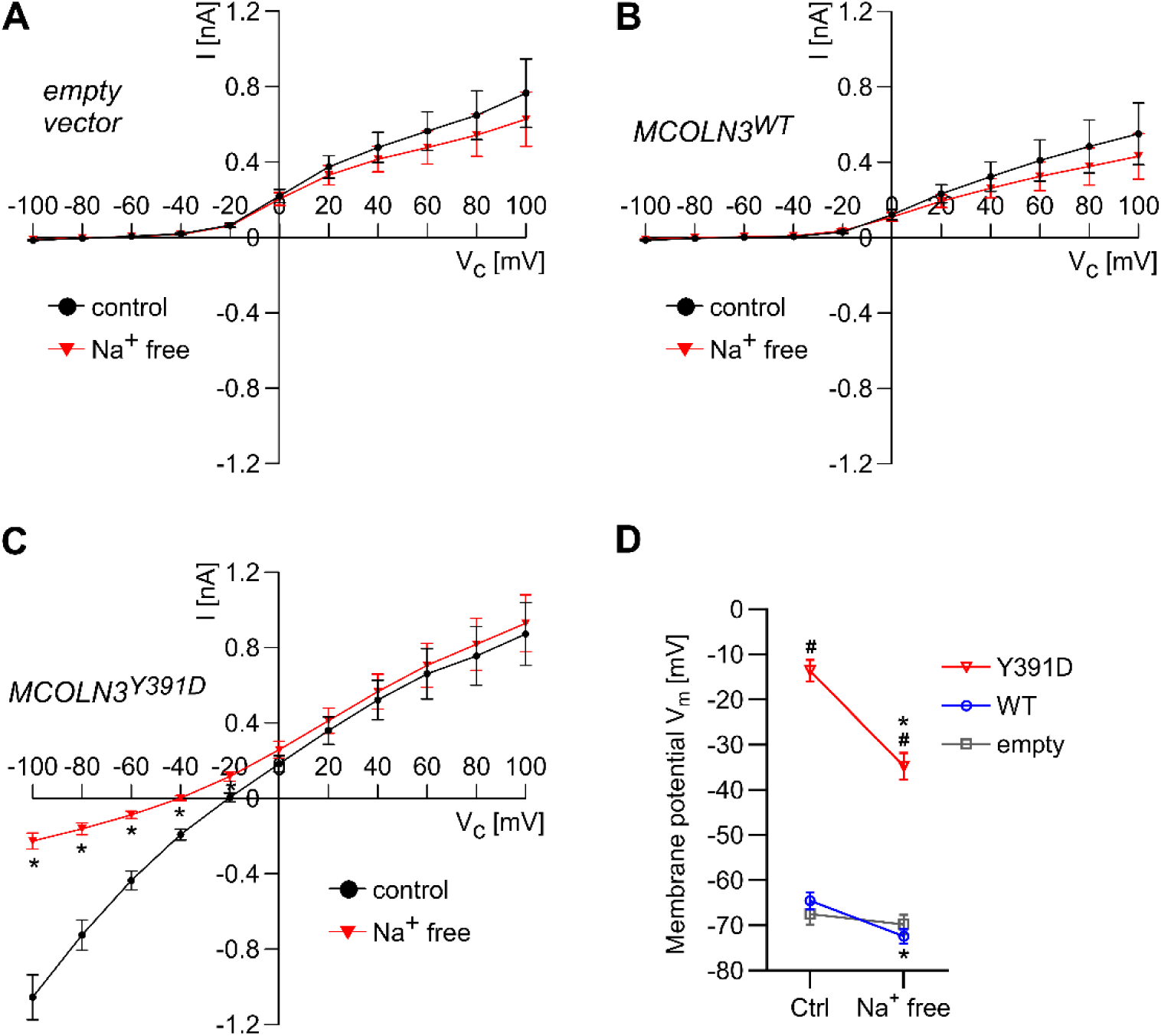
Whole-cell currents and membrane voltage in adrenal HAC15 cells transiently transfected with pIRES-CD8-plasmids containing wild type (*MCOLN3^WT^*) or mutant MCOLN3 (*MCOLN3^Y391D^*) under control condition and after replacement of extracellular sodium (Na^+^) by cell impermeable N-methyl-D-glucamine^+^ (NMDG^+^). Cell currents at different voltage-clamp steps (**A-C**) and cell membrane potentials (**D**) were measured using the whole-cell patch-clamp technique. (**A-C**) All cells showed outward currents typically present in adrenal cell lines and native adrenal glomerulosa cells. Overexpression of *MCOLN3^Y391D^* led to additional large inward currents at negative voltage-clamp steps (-100 mV up to -40 mV), which were absent in cells with empty vector and *MCOLN3^WT^* cells. Replacement of extracellular Na^+^ by NMDG^+^ (Na^+^ free, red triangles) led to a significant reduction of these inward currents in mutant cell, albeit not to the level of control cells. (**D**) Membrane voltage of cells with *MCOLN3^Y391D^* was depolarized compared to cells with empty vector or with *MCOLN3^WT^*, both under control and Na^+^ free conditions. Removal of Na^+^ led to a more hyperpolarized membrane potential in HAC15-*MCOLN3^WT^* and HAC15-*MCOLN3^Y391D^* cells, but to a higher extent in cells with HAC15-*MCOLN3^Y391D^* cells. Statistical significance (* and ^#^) was tested with the GraphPad Prism 10 software using a two-way ANOVA analysis with Tukey’s multiple comparison. \**P* < 0.05 for comparison of ctrl versus Na^+^ free solution in (A-D). #*P* < 0.05 for comparison of HAC15-*MCOLN3^Y391D^*cells vs HAC15-*MCOLN3^WT^* and empty control cells in (**D**); Data are presented as mean ±SEM (empty CD8-vector n=7, *MCOLN3^WT^*n=8, *MCOLN3^Y391D^* n=17). The number of experiments in (**A-C**) is equal to the number in (**D**) because the membrane voltage was measured alternately with the current measurements.

Next, we measured the cytosolic calcium activity in HAC15-*MCOLN3^Y391D^*cells under various extracellular conditions (Fig. 6B-C). These cells demonstrated significant elevation in basal intracellular calcium compared to the empty and HAC15-*MCOLN3^WT^* cells (*P*<0.05) (Fig. 6B). Cytosolic calcium activity was continuously monitored over time and upon exposure to extracellular solutions with varying calcium content. Cytosolic calcium levels in HAC15-*MCOLN3^Y391D^* cells showed a greater dependency on extracellular calcium levels than the HAC15-*MCOLN3^WT^*cells (Fig. 6C). This was indicated by the reduced intracellular calcium levels following exposure to Ca^2+^-free extracellular solution whereas high extracellular calcium levels caused a surge in cytosolic calcium. Furthermore, intracellular calcium levels were restored to baseline under control medium conditions. Cytosolic calcium levels in HAC15-*MCOLN3^WT^* and control cells were unaffected by altered calcium content in extracellular solutions (Fig. 6C). Overall, the fura-2 calcium measurements revealed an abnormal calcium influx and impaired calcium-handling capacity, which is likely caused by the abnormal basal activity of mutant *MCOLN3^Y391D^*and underlies the autonomous aldosterone production.

## Discussion

CYP11B2 expression, and therefore aldosterone production, is tightly regulated by intracellular calcium levels (43, 44). The somatic aldosterone-driver mutations identified in adrenal lesions in patients with primary aldosteronism often target mechanisms directly affecting intracellular calcium buffering, or indirectly, via altering the cell membrane potential triggering voltage-gated calcium channels to open. In this study, we report the first set of disease-causing mutations in *MCOLN3* identified in APAs from male patients with primary aldosteronism*. In vitro* investigations of the missense mutation showed increased aldosterone production by altering intracellular calcium flux.

Research into *MCOLN3* and its regulatory role in endocytic and autophagic pathways has increased the last few decades (24, 45–47). Subcellular locations of MCOLN3 include the plasma membrane, cytoplasm (28, 30), lysosomes, endosomes and autophagosome (24–26, 48–52). Human *MCOLN3* is located on chromosome 1 (1p22.3) and encodes for TRPML3 —a member of the mucolipin subfamily of transient receptor potential cation channels (TRPML) (53–55). TRPML3 consists of an ion-conducting pore domain formed by four subunits. Each subunit has six transmembrane segments and a pore loop located between segments 5 and 6. The pore loop is reported to consist of two helices (L444_I455 and M460_K466) located on either side of the ion selectivity motif (56). In mice, mutation in the residue at position 419 (in segment 5) located close to the ion transport region, leads to gain-of-function of the *MCOLN3* cation channel resulting in auditory disease (heterozygous) or death (homozygous) (26, 30–32, 47). The mouse and human *MCOLN3* share ∼91% sequence homology (57). While *MCOLN3* mutations have not been associated with disease in humans, the residues altered by *MCOLN3^Y391D^*and *MCOLN3^N411_V412delinsI^* identified in the APAs were found in close proximity to the ion transport motif (56).

In our study, *MCOLN3* was prominently expressed in the adrenal cortex and aldosterone-producing adrenal tumors. Our transcript data showing elevated *MCOLN3* expression in the adrenal cortex aligns with the study by Bergman *et al.* (2017) that indicated *MCOLN3* as one of thirty-seven genes with enriched expression in the normal human adrenal gland (specifically the cortex) compared to other human tissues (58). Interestingly, while MCOLN3 was abundantly expressed throughout the zona glomerulosa, its expression did not necessarily directly correlate with that of CYP11B2 as shown in adrenals with discontinuous zona glomerulosa CYP11B2 expression. This pattern of MCOLN3 and CYP11B2 expression in the normal adrenal suggests that these genes are regulated independently from one another.

While mutations in *MCOLN3* have been associated with disease in mice, to our knowledge, this study is the first report of disease-causing *MCOLN3* mutations in humans. In Varitint-waddler mice, the gain-of-function missense mutation A419P is located in segment 5 near the pore and causes deafness, circling behavior and abnormal pigmentation (31, 32, 59). This mutation results in a constitutively active inward current carried by calcium and sodium ions. (32, 60). Human and mice MCOLN3 protein sequence shares 96.2% similarity and 91.6% identity, which suggest that the mutations identified in the APAs might alter the protein function in a similar manner and cause abnormal ion transport. In this study, the human *MCOLN3* variants identified are both affected at locations close to that of the mouse A419P and the ion pore in segments 4 (*MCOLN3^Y391D^*) and 5 (*MCOLN3^N411_V412delinsI^*). Our studies focus on the *MCOLN3^Y391D^*variant that was identified in more than one APA. Overexpression of the *MCOLN3^WT^* variant in HAC15 cells did not alter transcription of *NR4A2* and *CYP11B2* nor the production of aldosterone. However, the single residue switch from tyrosine to aspartic acid at position 391 (*MCOLN3^Y391D^*) altered the steroidogenic enzyme expression and regulation of aldosterone production. The fact that *MCOLN3* expression is enriched in the adrenal cortex but that overexpression does not significantly alter steroidogenesis in these cells suggests that *MCOLN3^WT^* is not directly involved in the regulation of aldosterone synthesis. The physiological function of *MCOLN3^WT^*in the adrenal cortex remains to be elucidated.

Co-expression of CYP11B2 and CYP17A1 is characteristic of adrenal tumors harboring *KCNJ5*-(40, 61) and *SLC30A1-* mutations (17). Consequently, elevated biosynthesis of 18-hydroxycortisol and 18-oxocortisol is detected in these patients and considered biomarkers that are utilized in the diagnostic approach for primary aldosteronism (37, 62–65). In our study, CYP17A1 tumor immunoreactivity accompanied that of CYP11B2 and MCOLN3. Due to the the lack of patient serum availability, we were unable to determine whether the hydrid steroids are increased in these patients. However, elevated hybrid steroids measured in the adrenocortical cell experimental medium suggest that these should not be ruled out in future investigations. The modest increase in *CYP17A1* mRNA in HAC15-*MCOLN3^Y391D^* cells could be attributed to the fact that CYP17A1 expression is already high in this adrenal cell line, limiting the possibility for further induction. Nevertheless, our cell experiments showed that the levels of transcripts and steroids involved in the production of aldosterone was significantly induced in the HAC15-*MCOLN3^Y391D^*cells *vs* the HAC15-*MCOLN3^WT^* cells. The induction of transcripts and steroid metabolites associated with the zona fasciculata was less prominent or unaffected. The data suggest that the *MCOLN3* mutations targeted aldosterone biosynthesis more specifically and were not responsible for an overall induction of steroidogenesis. This further implicates these *MCOLN3* variants as a disease-causing mutation in patients with primary aldosteronism. It was interesting to note that the immunoreactivity of MCOLN3 and CYP11B2 was more profound in the tumors harboring *MCOLN3^Y391D^* than the tumor harboring *MCOLN3^N411_V412delinsI^.* The *MCOLN3^N411_V412delinsI^*variant was identified in an African American patient who, out of the three, was the youngest and presented with more severe hypertension. Previous studies have found that patients of African heritage had a positive correlation between the severity of hyperaldosteronism and blood pressure (66, 67) and a higher susceptible to sodium-sensitive hypertension and hyperaldosteronism (33–35). These factors increases the risk of congestive heart failure, end-stage renal disease and atherosclerosis in this demographic (68).

Investigations into the functional consequences of *MCOLN3^Y391D^*on the cellular electrophysiological properties demonstrated an increase in depolarization of the membrane potential in HAC15-*MCOLN3^Y391D^*cells compared to the control and HAC15-*MCOLN3^WT^* cells. The impact of extracellular calcium levels on the cytosolic calcium influx observed in the HAC15-*MCOLN3^Y391D^* cells suggests that this could result in gain-of-function leading to a constitutively active calcium-permeable channel phenotype. On the other hand, a layer of complexity is added by the data showing the impact of extracellular sodium on cell membrane potential and calcium influx. Increased sodium conductance, and therefore sodium-induced depolarization, has been associated with other aldosterone-driver mutations (7). As a known cation-permeable channel, it is possible that *MCOLN3^Y391D^*-induced calcium influx occurs directly as well as indirectly via sodium-induced cell membrane depolarization resulting in a cumulative increase in cytosolic calcium levels (Fig. 8).

**Figure 8.**
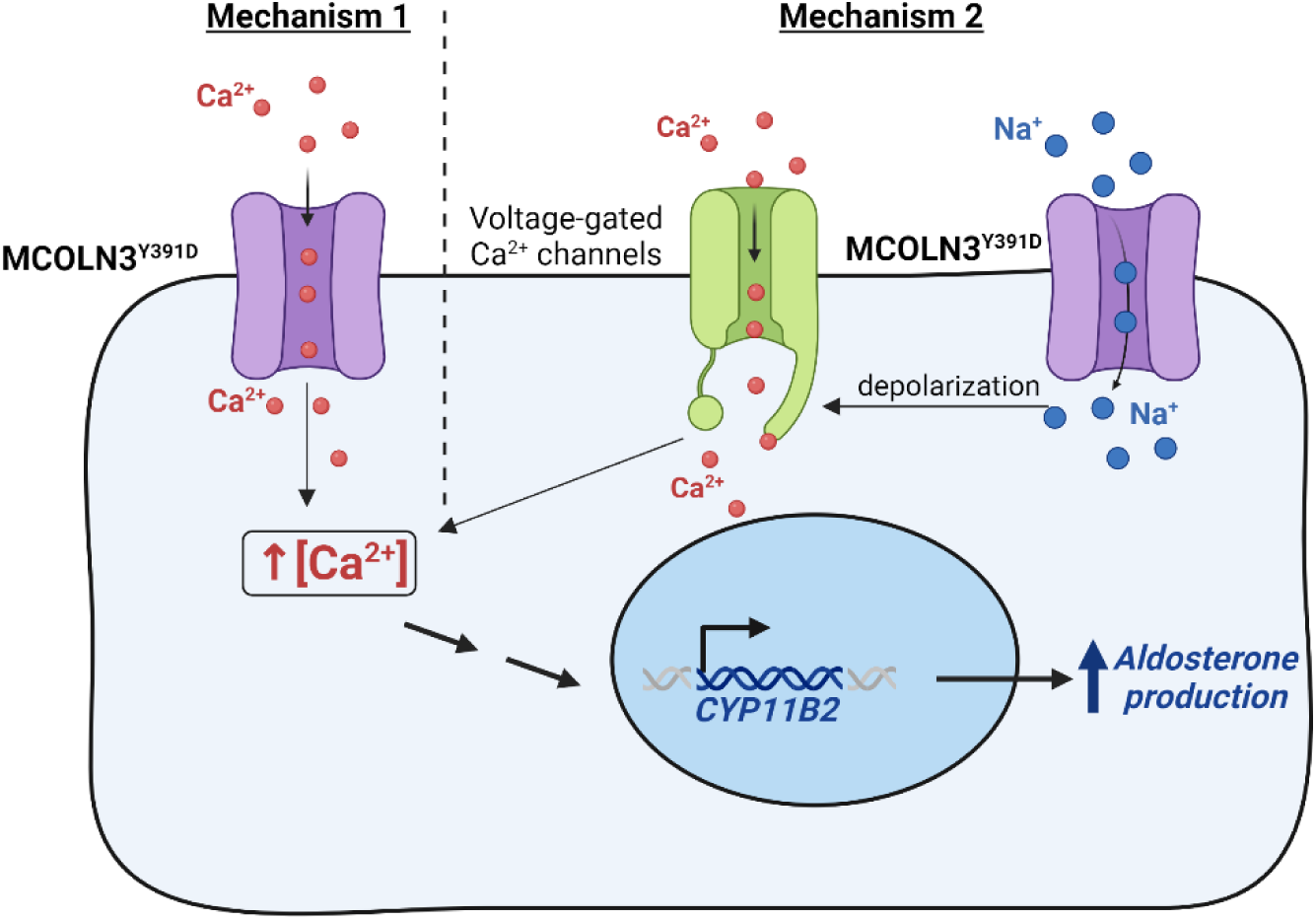
Mechanisms of cytosolic calcium (Ca^2+^) influx caused by *MCOLN3^Y391D^*. Mechanism 1 proposes that *MCOLN3^Y391D^* resulted in a gain-of-function calcium permeable channel which directly increases cytosolic calcium levels. In mechanism 2, *MCOLN3^Y391D^* altered the ion selectivity of *MCOLN3* resulting in the influx of sodium ions (Na^+^) which causes cell membrane depolarization and the activation of voltage-gated calcium channels. Elevated cytosolic calcium levels induce the transcription of aldosterone synthase (CYP11B2) and hyperaldosteronism. Image created with BioRender.com

While the exact mechanism of *MCOLN3^Y391D^*-induced elevated CYP11B2 expression and aldosterone production remains to be elucidated, it is evident that adrenal cells expressing *MCOLN3^Y391D^* exhibit an increased influx of calcium. The data presented in this study implicate *MCOLN3^Y391D^* in the pathogenesis of primary aldosteronism. Furthermore, this study is the first to report *MCOLN3* mutations in APAs and to associate mutations in *MCOLN3* with disease in humans.

## Materials and Methods

### Patients and archival FFPE APA and adrenal tissue

Consenting patients from the University of Michigan, University of Pennsylvania, and Vanderbilt University with a confirmed unilateral primary aldosteronism diagnosis (based on the institutional consensus or Endocrine Society’s clinical practice guidelines) were included in this study. The selection criteria further considered the availability of FFPE blocks of resected adrenals. Baseline clinical information was collected from each patient. Adrenal tissue from healthy subjects was obtained from the University of Michigan kidney transplant donor program. This study was conducted with the approval of the institutional review boards of the University of Michigan (HUM00083056 and HUM00069665), University of Pennsylvania (IRB# 831990), and Vanderbilt University Medical Center (IRB# 150995). We will share limited NGS data on request.

### Immunohistochemistry of FFPE adrenal tissue

5 µm thick serial sections of the FFPE APA and adrenal tissue were prepared, and the first few sections used for hematoxylin and eosin (H&E)- and IHC as previously described (17, 69, 70). In brief, FFPE tissue sections were deparaffinized and subsequently heated for 15 min in the appropriate antigen retrieval solution (see table 1 for antibody-specific detail). Non-specific binding was limited by incubating samples for 10 min with peroxidase and a further 1 h with 10 % goat serum prepared in Triton X-100 phosphate-buffered saline. Next, tissue sections were incubated for 1 h at room temperature with the specific primary antibody after which detection was achieved using the Polink-2 horseradish peroxidase (HRP) Plus Mouse DAB System (GBI Labs). Lastly, slides were counterstained with Harris hematoxylin, dehydrated and cover slipped prior to being scanned using the PathScan Enabler IV (Meyer Instruments).

### CYP11B2 IHC-guided isolation of genetic material from FFPE adrenal tissue

The first two slides of the serial sections were used for H&E and CYP11B2 IHC. The CYP11B2-stained section was then used as a reference to dissect the tentative CYP11B2-positive tumor region from the subsequent 6-8 unstained sections using a scalpel under an Olympus SZ-40 microscope. Adjacent adrenal tissue regions were also collected from the unstained FFPE slides. The Qiagen AllPrep DNA/RNA FFPE kit was used to isolate the genomic DNA and RNA that was then used in downstream processes.

### Whole exome sequencing (WES)

Genomic DNA isolated from CYP11B2 IHC-guided APAs that were found to be negative for all known aldosterone-driver mutations, were analyzed with WES (14, 17, 70). Matched-adjacent adrenal gDNA was used for germline variant identification. The Clinical Laboratory Improvement Amendments-compliant sequencing lab performed all WES analyses using standard protocols. In brief, gDNA (500 ng) was fragmented to 250 bp peak target size fragments using a Covaris S2 ultrasonicator and concentrated using AMPure beads. Next, the fragments were subjected to end-repair, A-base addition, Illumina indexed adapter ligation, and size selection on a NuSieve 3 % agarose gel (Lonzo). 1 µg of the library, prepared through the amplification and purification of 300-350 bp fragments using Illumina index primers and AMPure beads, respectively —was hybridized to the Agilent SureSelect Human All Exon v.4. The capture and enrichment of targeted exon fragments were carried out as per the manufacturer’s recommendations. Agilent 2100 Bioanalyzer and DNA 1000 reagents were used to analyze the paired-end whole-exome libraries prior to sequencing using the Illumina HiSeq 2500 sequencing system (Illumina). Primary base call files were converted into FASTQ sequence files using the bcl2fastq converter tool bcl2fastq-1.8.4 in the CASAVA 1.8 pipeline.

### Targeted gene sequencing using Ion Torrent next-generation sequencing (NGS) and Sanger sequencing

The WES finding of *MCOLN3* variants were confirmed using targeted NGS and Sanger sequencing of gDNA from the APAs identified with *MCOLN3* mutation and their matched-adjacent tissue. For confirmation via targeted Ion Torrent-based NGS, we developed a custom AmpliSeq multiplex PCR-based NGS panel that targets the complete coding sequencing of known mutated genes. Genes included in this panel (v7), include: *ATP1A1*, *ATP2B3*, *CACNA1D*, *CACNA1H*, *CADM1*, *CLCN2*, *CTNNB1*, *GNA11*, *GNAQ*, *KCNJ5*, *MCOLN3*, and *SLC30A1*. Briefly, barcoded NGS libraries were constructed from up to 20 ng of isolated gDNA using the Ion AmpliSeq Library Kit Plus (Thermo Fisher Scientific, Waltham, MA) prior to pooling for template preparation and sequencing on a Ion Chef Instrument and Ion GeneStudio S5 Prime NGS System, respectively. Raw NGS data was processed and aligned to the human genome (hg19) using Ion Torrent Suite, and after sample-level quality control (QC) filtering, variants were identified and annotated using the Ion Torrent Suite variantCaller plugin and ANNOVAR, respectively. Annotated variants were manually filtered to remove germline variants and sequencing or FFPE artifacts by an experienced molecular pathologist (A.M.U.), and final prioritized variants were visually confirmed using the Integrative Genomics Viewer (IGV; Broad Institute, Cambridge, MA). Finally, for Sanger validation, PCR amplification of gDNA was achieved following the instruction of the Promega GoTaq Flexi DNA Polymerase using 35 cycles and an annealing temperature of 56 °C. The specific *MCOLN3* primers used for sequencing of PCR products are provided in Table 2. Sanger chromatograms were analyzed using SeqMan Ultra software (V17.3.1) from DNASTAR, Inc.

**Table 2.** Immunohistochemistry staining detail.

| Target | Antigen retrieval solution<br>(*Vector Laboratories Inc.*) | Primary antibody detail |
| --- | --- | --- |
| Human CYP11B2 | Tris/EDTA buffer (pH 9) | Mouse monoclonal, clone 41-17B (1:1250),<br>(Millipore Cat# MABS1251,<br>RRID:AB_2783793) |
| Human CYP17A1 | Tris/EDTA buffer (pH 9) | Rabbit polyclonal (1:2000),<br>(LSBio (LifeSpan) Cat# LS-B14227,<br>RRID:AB_2857939) |
| Human MCOLN3 | Sodium citrate buffer (pH 6) | Rabbit polyclonal (1:750),<br>Sigma-Aldrich Cat# HPA018106,<br>RRID:AB_1853647) |

**Table 3.**
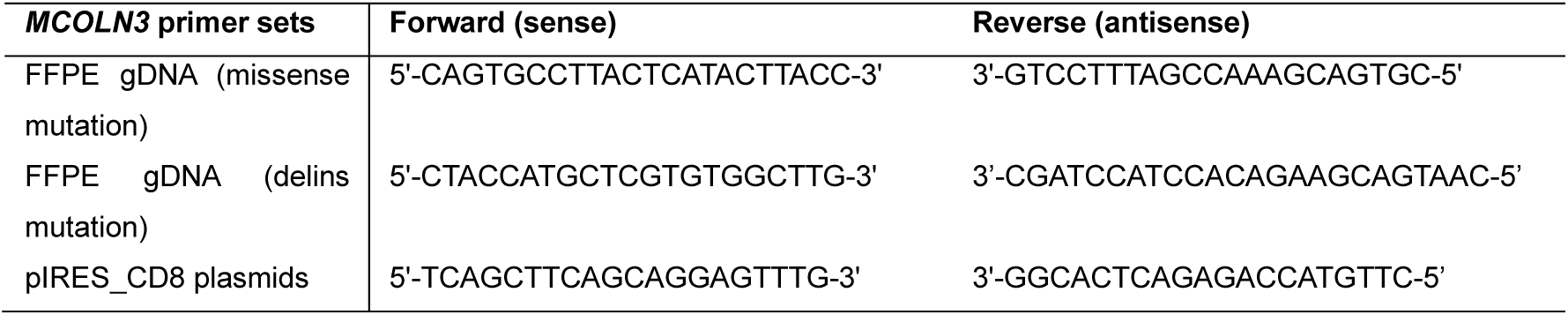
Primer set sequences for human *MCOLN3*.

### Expression vectors

GenScript supplied the cDNA open reading frames (ORF) of human wild type *MCOLN3* gene (OHu23350) and *MCOLN3* p.Y391D variant (obtained through site-directed mutagenesis of the wild type clone) in the pcDNA3.1-C-(k)DYK backbone. The wild type *MCOLN3* and *MCOLN3* p.Y391D plasmid DNA was cloned in the pIRES-CD8 vector and used in all adrenocortical cell line experiments.

### Cell culture and reagents

Human adrenocortical HAC15 cell line was routinely cultured in growth media consisting of Dulbecco’s modified Eagle’s medium (DMEM)-F12 (Gibco) supplemented with 10 % cosmic calf serum (Hyclone), 1 % ITS (insulin-transferrin-selenium; Corning), and antibiotics, 1 % penicillin/streptomycin (Gibco) and 0.1 % gentamicin (Gibco). Cells were grown at 37 °C in a humidified atmosphere of 5 % CO2.

### Development of HAC15 cells expressing *MCOLN3* variants

For the generation of HAC15 cells expressing variants of *MCOLN3*, HAC15 cells (2x10^6^ live cells) were transfected with 1 µg of plasmid DNA via electroporation using an AMAXA nucleofector kit following the manufacturer’s recommendation (Lonza Group Ltd., Basel, Switzerland). Plasmids included the empty pIRES-CD8 plasmid and pIRES-CD8 plasmids containing wild type *MCOLN3* (*MCOLN3^WT^*) or *MCOLN3^Y391D^*.

In brief, HAC15 cells were cultured to 70-80 % confluency in growth medium before use. Cells were trypsinized (3 mL) and trypsin was inactivated by the addition of 10 mL growth media. Cell concentration was determined and the calculated suspension containing *2 million live cells* X *number of transfections*, was transferred to a 15 mL conical tube. Cells were pelleted by centrifugation (4 min at 1000 rpm) and all traces of serum, which could hinder transfection efficiency, was removed by washing the isolated cell pellet with 3 mL BSA (0.5 % in 1X PBS) followed again by centrifugation. The serum-free cell pellet was carefully resuspended in supplemented Nucleofector solution (*100 µL per 2 million cells* X *number of transfections*) and the time noted as to not exceed the 15 min cells-to-solution exposure, as per the manufacturer’s instructions (Lonza).

The experiment proceeded with single plasmid transfections: 100 µL of the cell suspension in Nucleofector solution (NCS) was carefully mixed with 1 µg plasmid DNA aliquot (in 5 µL) using a P1000 pipette. The cell solution was then transferred to an AMAXA certified cuvette (provided in kit) and subjected to electroporation using the T20 Nucleofector® program. Immediately post-electroporation, 500 µL warm growth medium was added to the cuvette for cell recovery. Using a plastic pipette, the transfected cells were gently divided, in a dropwise-manner, between 3 wells of a 12-well plate containing 600 µL pre-warmed growth media. Cells were incubated for 48 h after which media was collected and stored at -20 °C and plates stored at -80 °C for future protein quantification or RNA isolation.

### Total protein extraction and quantification

Cells were lysed in 400 µL of mammalian protein extraction reagent (M-PER) for 5 min with agitation and centrifuged for 10 min at 10,000 *x g.* The total protein content was quantified using the bicinchoninic acid (BCA) protein assay (Pierce Chemical Co.).

### Aldosterone ELISA assay

An in-house competitive enzyme-linked immunosorbent assay (ELISA) was used to quantify aldosterone levels in the experimental cell medium (71, 72). Flat bottom, high-binding 96-well plates (Fisher Scientific) were coated with 2 µg of goat anti-mouse IgG (Fc specific, Sigma-Aldrich; RRID: AB_260486) prepared in 100 µL coating buffer (BupH™ Carbonate-Bicarbonate Buffer, pH 9.4-9.6) (Pierce) and incubated overnight at 4 °C. The plates were washed with 4 x 300 µL of the wash buffer (phosphate-buffered saline (PBS) containing 0.1% Tween) prior to being coated with an aldosterone antibody (Aldo A2E11-6G1; 1:2500, RRID: AB_2892670). 50 µL of the cell medium samples and aldosterone standards (prepared in experimental medium) were pipetted into the respective wells and incubated on a shaker with 50 µL HRP-conjugated aldosterone (Aldo-3CMO-HRP; 1:20000) for 2h. The wells were then washed, the substrate solution containing tetramethylbenzidine (100 µL; Fisher Scientific) added and incubated on a shaker for a further 20-30 min. Finally, the reaction was stopped with the addition of 50 µL of sulfuric acid (1N) and the absorbance measured within 15 min at 450 nm using the Biotek Epoch Gen5 spectrophotometer (Agilent). The unknown aldosterone concentrations were extrapolating from the standard curve generated by the Gen5 software (v.2.09). Aldosterone concentrations were normalized to total protein and expressed as ng/mg total protein.

### RNA isolation from cell experiments

Total RNA was isolated from cells using the RNeasy Plus Mini kit (Qiagen) and stored at -80 °C. Prior to use in downstream processes, RNA was quantified, and RNA purity and integrity inspected using the Nano Drop spectrophotometer instrument (Nano Drop Technologies).

### Reverse transcription and quantitative real-time PCR (qRT-PCR)

RNA was reverse transcribed using the High-Capacity cDNA Reverse Transcription Kit (Applied Biosystems). Real-time quantitative PCR (qRT-PCR) analyses were performed using 5 ng and 25 ng of cDNA from snap frozen tissue and cell experiments, respectively, and Taqman Fast Universal Master Mix (Life Technologies) as per manufacturer’s recommendations and analyzed using the StepOne Plus Fast Real-Time PCR system (Applied Biosystems). The primer-probe sets of human transcripts purchased from Life Technologies include peptidylprolyl isomerase A (*PPIA*, cyclophilin A; Hs99999904_m1), NURR1 (*NR4A2*; Hs00428691_m1), and mucolipin 3 (*MCOLN3*; Hs00539554_m1).

*CYP17A1, CYP11B1,* and *CYP11B2* primer-probe set was designed in-house and purchased from Integrated DNA Technologies (73, 74). Target gene expression was normalized to the expression of *PPIA* (housekeeping gene) and quantified using the comparative threshold cycle method using StepOne software (v.2.3). Transcript data was expressed relative to the empty pIRES-CD8 control.

### Whole-cell patch-clamp experiments and calcium measurement assays

HAC15 cells were cultured as previously described. For electrophysiologic investigation, HAC15 cells, seeded on fibronectin/collagen-coated glass cover slips (Corning Life Sciences, Amsterdam, the Netherlands), were transfected with 1 μg of empty pIRES-CD8 plasmid, or with 0.5 µg of pIRES-CD8 plasmids containing *MCOLN3^WT^*or *MCOLN3^Y391D^* using Lipofectamine 3000. For the *MCOLN3*-transfections, plasmid concentration was adjusted to 1 μg with empty pIRES-CD8 to ensure sufficient bead binding. Experiments were performed 24-30 h after transfection. Transfected cells were identified by binding anti-CD8 coated Dynabeads (LifeTechnologies GmbH, Darmstadt, Germany) and used for whole-cell patch-clamp experiments and cytosolic calcium measurements. Whole-cell patch-clamp recordings were performed at room temperature using an EPC 10 amplifier (Heka, Lambrecht, Germany) and a Powerlab Data Acquisition System (ADInstruments GmbH, Spechbach, Germany). The PatchMaster v2x50- and FitMaster v2x90.5 software (Heka, Lambrecht, Germany) and the LabChartPro v7 software (ADInstruments GmbH, Spechbach, Germany) were used for data acquisition and analysis. Cell membrane potential was measured in current clamp mode with subtraction of 10 mV liquid junction potential. Inward and outward currents were measured in voltage clamp mode by a series of clamp steps (from -100 to +100 mV incremented in 20 mV steps). Patch pipettes with 5-10 MΩ were used for the recordings. The patch pipette solution contained (mM): 95 K-gluconate, 30 KCl, 4.8 Na2HPO4, 1.2 NaH2PO4, 5 glucose, 2.38 MgCl2, 0.726 CaCl2, 1 EGTA, 3 ATP, pH 7.2. The extracellular Ringer-type control solution contained (mM): 145 NaCl, 3.6 KCl, 5 glucose, 1 MgCl2, 1.3 CaCl2, 10 HEPES, pH 7.4. For Na^+^-free solution, Na^+^ was replaced by N-methyl-D-glucamine (NMDG^+^).

For cytosolic calcium activity assays, transfected HAC15 cells were loaded with 5 µM Fura 2-AM solution (+ power load) for 1h 15 min. Next, cytosolic calcium activity was measured at basal conditions as well as after treatment with extracellular solutions with various calcium content: control Ringer solution (1.3 mM Ca^2+^; ctrl); 5 mM (high-Ca^2+^); and Ca^2+^-free solution (calcium was omitted when weighing the substances, but possible traces of calcium were not removed by complexation with EGTA in order to prevent the cells from detaching). Fura-2-fluoresence intensity was measured at 510 nm after excitation at 340 nm and 380 nm. Fura-2 ratio data represents the mean ratio ±SEM of *n* dishes, each from which the average signal from *X* number of confirmed transfected cells per dish, were calculated (empty vector, n=10; with *MCOLN3^WT^*, n=11; with *MCOLN3^Y391D^*, n=10). Signals from untransfected cells of the matched experimental dishes served as internal controls.

### Statistical Analysis

All cell experiment results are presented as the mean ±SEM of at least four independent experiments performed in triplicate. Statistical analyses were performed using an ordinary one-way ANOVA with a Tukey’s multiple comparison test. Snap frozen tissue mRNA data is presented as the mean ±SD or median with 95 % CI of at least four samples and analyzed using a Kruskal-Wallis with a Dunn’s multiple comparison test. Statistical analyses were performed using GraphPad Prism version 10 (GraphPad Software, Inc., San Diego, CA). * *P*<0.05; ** *P*<0.01; *** *P*<0.001; \*\*\*\**P*<0.0001.

## Supporting information

Manuscript file

## Acknowledgments

This work was supported, in part, by grants from: the Michigan Biologic Research Initiative for Sex Differences in Cardiovascular Disease (M-BRISC) program at the Frankel Cardiovascular Center, University of Michigan and the National Institutes of Health Award R01 DK106618 (**for WER**); the American Heart Association Award (23POST1020221) (**for DVR**); National Heart, Lung, and Blood Institute (NHLBI) (R01HL174676), National Center for Advancing Translational Sciences (NCATS)/Michigan Institute for Clinical and Health Research (UL1TR002240), the University of Michigan Frankel Cardiovascular Center McKay Grant Award (G028398), University of Michigan Pandemic Research Recovery Grant and the American Heart Association (20CDA35320016) (**for JR**); R01DK081662 (**for JML**), NCATS (UL1TR002243 to the Vanderbilt Clinical Translational Science Award program), NCI (2P30CA068485 to the Vanderbilt Translational Pathology Shared Resource); National Cancer Institutes of the National Institutes of Health (K08 CA270385) (**for HW**). The plasmid DNA preparation was performed at the University of Michigan Vector Core and was supported by a grant from the National Institute of Diabetes and Digestive and Kidney Diseases (NIDDK) to the University of Michigan Center for Gastrointestinal Research (5P30DK034933).

