## Supplementary material for "Somatic Mutations in *MCOLN3* in Aldosterone-Producing Adenomas cause Primary Aldosteronism": Manuscript file

### Supplemental file

LC-MS/MS analyses of steroid levels in cell culture medium.

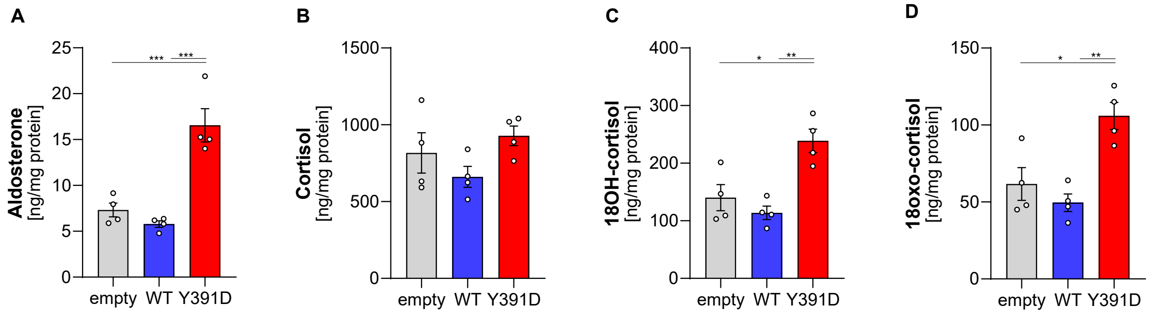


**Supplemental Fig. 1|** LC-MS/MS analysis of aldosterone, cortisol, 18-hydroxycortisol (18OH-cortisol) and 18-oxocortisol measured in cell experiment medium.

#### Supplemental Table 1 | Selection of somatic variants detected by whole exome sequencing

| Sample | Gene | Chromosome position | Amino acid change | Mutation type | Variant Allele Frequency in tumor (VAF) |
|---|---|---|---|---|---|
| **UM102** | *CELA3B* | chr1:22307199 | | exon_loss | 7.00% |
| | *GRK3* | chr22:26110531 | p.Glu550Lys | missense | 7.00% |
| | *KRTAP4-7* | chr17:39240790 | p.Arg111His | missense | 24.00% |
| | *KRTAP9-6* | chr17:39421886 | p.Tyr86Cys | missense | 10.00% |
| | *MARS* | chr12:57905563 | p.Ser484Cys | missense | 7.00% |
| | ***MCOLN3*** | **chr1:85488008** | **p.Tyr391Asp** | **missense** | **36.00%** |
| | *OR7E24* | chr19:9361740 | p.Phe11fs | frameshift | 19.00% |
| | *SCN1B* | chr19:35524610 | p.Val139Ile | missense | 12.00% |
| | *SFMBT2`* | chr10:7326093 | p.Pro182His | missense | 12.00% |
| | *UGT3A1* | chr5:35954326 | p.Ala517Val | missense | 7.00% |
| **ELA25** | ***MCOLN3*** | **chr1:85022325** | **p.Tyr335Asp** | **missense** | **33.70%** |
| | *PLEKHG4B* | chr5:156173 | p.Arg415Trp | missense | 30.30% |
| | *CDH10* | chr5:24487719 | p.Arg771Trp | missense | 30.20% |
| | *MNT* | chr17:2387260 | p.Gly464Ser | missense | 30.20% |
| | *ZZEF1* | chr17:4104662 | p.Ser515Leu | missense | 29.70% |
| | *TEP1* | chr14:20408112 | p.Arg110Trp | missense | 25.80% |
| | *SLC6A13* | chr12:237241 | p.Arg205Gly | missense | 25.30% |
| | *TNKS1BP1* | chr11:57309068 | p.Glu1215Lys | missense | 21.00% |
| | *CPLANE1* | chr5:37195954 | p.Gln1239Ter | stopgain | 20.60% |
| | *YTHDC2* | chr5:113563868 | p.Ala818Thr | missense | 19.10% |
| | *ZNF462* | chr9:106927310 | p.Tyr1133Phe | missense | 18.90% |
| | *ATP8B1* | chr18:57731798 | p.Glu4Lys | missense | 17.50% |
| | *VPS13B* | chr8:99431619 | p.Glu1055Asp | missense | 17.30% |
| | *SPPL2C* | chr17:45845979 | p.Ser358Phe | missense | 16.00% |
| | *USP6NL* | chr10:11462770 | p.Gly737Arg | missense | 15.90% |
| | *NTRK2* | chr9:85021361 | p.Lys814Met | missense | 15.80% |
| | *FAT3* | chr11:92798088 | p.Leu1692Gln | missense | 15.80% |
| | *RNF19A* | chr8:100258600 | p.His825ThrfsTer13 | frameshift | 15.20% |

| | | | | | |
|---|---|---|---|---|---|
| | *PCDHGC4* | chr5:141486710 | p.Gln513Ter | stopgain | 15.10% |
| | *ZSCAN32* | chr16:3382965 | p.Tyr449Asn | missense | 15.00% |
| | *GRXCR2* | chr5:145872794 | p.Glu59Lys | missense | 15.00% |
| | *ACADM* | chr1:75732897 | p.Met87Ile | missense | 14.90% |
| | *VSIG10L2* | chr11:125955822 | p.Glu738Lys | missense | 14.50% |
| | *TESK1* | chr9:35606026 | p.Asn88Tyr | missense | 14.10% |
| | *SPHKAP* | chr2:228018185 | p.Ala890Val | missense | 13.90% |
| | *ZNF483* | chr9:111541959 | p.His342Tyr | missense | 13.50% |
| | *FRY* | chr13:32147895 | p.Pro447Leu | missense | 13.20% |
| | *MPP5* | chr14:67292678 | p.Pro179Ser | missense | 13.10% |
| | *KMT5B* | chr11:68171043 | p.Glu317Lys | missense | 12.80% |
| | *GTF2H1* | chr11:18347990 | p.Cys171Tyr | missense | 12.80% |
| | *LPCAT3* | chr12:6977462 | p.Pro418Ser | missense | 12.70% |
| | *ICAM5* | chr19:10292044 | p.Pro228Leu | missense | 12.50% |
| | *NFAT5* | chr16:69693986 | p.Glu1387Asp | missense | 12.40% |
| | *ICE1* | chr5:5463352 | p.Glu1340Lys | missense | 12.30% |
| | *EXTL3* | chr8:28717235 | p.Asp392Glu | missense | 12.10% |
| | *ZSWIM1* | chr20:45883723 | p.Gln377His | missense | 12.10% |
| | *C4BPA* | chr1:207124270 | p.Asp204Asn | missense | 12.00% |
| | *KAT6A* | chr8:41934279 | p.His1314Arg | missense | 11.90% |
| | *SPATA13* | chr13:24297583 | p.Asp463Ala | missense | 11.90% |
| | *DOT1L* | chr19:2191196 | p.Glu150Val | missense | 11.60% |
| | *RBCK1* | chr20:427323 | p.Pro305Leu | missense | 11.50% |
| | *SCLY* | chr2:238082054 | p.Glu208Lys | missense | 11.30% |
| | *LRRC8D* | chr1:89933384 | p.Thr106Ala | missense | 10.60% |
| | *AXIN2* | chr17:65557833 | p.Ser263Tyr | missense | 10.60% |
| | *TUBAL3* | chr10:5393792 | p.Thr356Ala | missense | 10.40% |
| | *TMTC3* | chr12:88153398 | p.His100MetfsTer6 | frameshift | 9.60% |
| | *SRRM2* | chr16:2766205 | p.Ser1893Pro | missense | 9.10% |
| | *GJB5* | chr1:34757509 | p.Cys60Tyr | missense | 8.90% |
| | *SYDE1* | chr19:15113844 | p.Gly697Ser | missense | 6.50% |
| | *ZNF559* | chr19:9341912 | p.Thr154Lys | missense | 6.30% |

### Supplemental Table 2 | Results of targeted next-generation sequencing

| Sample | *Gene* | Nucleotide Change | Amino Acid Change | Flow-corrected Read Depth (FDP) | Variant Allele Frequency (VAF) |
|---|---|---|---|---|---|
| **UM102** | *MCOLN3* | c.T1171G | p.Y391D | 1998 | 35.9 |
| **ELA25** | *MCOLN3* | c.T1171G | p.Y391D | 2000 | 33.9 |
| **AA30** | *MCOLN3* | c.1232_1234del | p.N411_V412delinsI | 1983 | 36.1 |
